# Cell-material interactions in 3D bioprinted plant cells

**DOI:** 10.1101/2024.01.30.578043

**Authors:** Imani Madison, Maimouna Tahir, Lisa Van den Broeck, Linh Phan, Timothy Horn, Rosangela Sozzani

## Abstract

3D bioprinting is an additive manufacturing technology with promise towards facilitating tissue engineering and single-cell investigations of cellular development and microenvironment responses. 3D bioprinting is still a new technology in the field of plant biology so its optimization with plant cells is still widely needed. Here, we present a study in which 3D bioprinting parameters, such as needle gauge, extrusion pressure, and scaffold type, were all tested in 3D bioprinted Tobacco BY-2 cells to evaluate how cell viability is responsive to each parameter. As a result, this study revealed an optimal range of extrusion pressures and needle gauges that resulted in an optimum cell viability. Furthermore, this study applied the identified optimal 3D bioprinting parameters to a different cell line, *Arabidopsis* root protoplasts, and stress condition, phosphate starvation, to confirm that the identified parameters were optimal in a different species, cell type, and cellular microenvironment. This suggested that phosphate-starved bioprinted *Arabidopsis* cells were less viable by 7 days, which was consistent with whole root phosphate starvation responses. As a result, the 3D bioprinter optimization yielded optimal cell viabilities in both BY-2 and Arabidopsis cells and facilitated an applied investigation into phosphate starvation stress.

## Introduction

3D bioprinting is an additive manufacturing (AM) technique that employs “bottom-up” layering of biological materials to form 3D geometric models. 3D bioprinting has developed rapidly in the past few decades towards facilitating tissue engineering and other applications (Bhargava et al., 2022; Jadhav and Jadhav, 2022; Jose et al., 2016). Tissue engineering by way of 3D bioprinting seeks to form distinct structures from viable plant or animal cells based on a particular programmable biomatter deposition (Bhargava et al., 2022; Jose et al., 2016; Mahmood et al., 2022; Jadhav and Jadhav, 2022). More generally, 3D bioprinting transfers biological materials onto a substrate in a programmed manner to accomplish a biological function (Jose et al., 2016; Mahmood et al., 2022). Extrusion-based 3D bioprinting is a subset of 3D bioprinting that transfers material, or bioink, by extruding it from the bioprinter nozzle (Jose et al., 2016). In plant tissue culture, 3D bioprinting can be used to replicate internal and external plant cells environments using hydrogel encapsulation materials that comprise the bioink (Jose et al., 2016; Mahmood et al., 2022; Ngo et al., 2018). By modeling the interactions between cells and their environment, scientists can begin to observe mechanical properties of plant development at a cellular level (Van den Broeck et al., 2022; Mehrotra et al., 2020; Hamant and Moulia, 2016; Miyashima et al., 2019). A 3D bioprinter can employ plant cell encapsulation in various mechanisms. However, these mechanisms should be chosen carefully based on the desired application of the bioprinted material, the properties of the cell-line being used, and the ideal characteristics for bioink material compatibility (Van den Broeck et al., 2022; Mahmood et al., 2022; Jose et al., 2016).

The Tobacco Bright Yellow 2 (BY-2) cell line, derived from a Nicotiana Tabacum (NT-1) cell suspension, is a cell culture commonly used for research surrounding the biological systems of plants due to its uniquely high uniformity and proliferation rate in optimal medium conditions (Nagata et al., 1992, Santos et al., 2016). The simplicity, and therefore predictability, of BY-2 cells serves as an opportunity for understanding more complex systems of plants at a cellular level.

The homogeneity of this cell line enables scientists to apply observed effects of various stimuli at a cellular level to a more complex multicellular tissue using a computer-aided analysis that generates a statistical correlation of cellular interactions to predict the behavior of more complex systems (Santos et al., 2016; Calcutt et al., 2021; Jose et al., 2016). Moreover, the characteristics of the tobacco BY-2 cell line, such as its high rate of division, synchronicity, high susceptibility to gene transformations, and efficiently large biomass potential, make it a particularly ideal experimental material for a wide range of parameter optimization applications in plant biology because they ensure reproducibility and consistent points of reference for studies examining cell cycle specific events (Santos et al., 2016; Calcutt et al., 2021; Nagata et al., 1992).

When engineering tissues for cell model investigations, it is important to evaluate the parameters of the physical and physiological environment that are responsible for the viability of the cell line *in vivo*. Many of these environmental parameters can be redesigned through the integration of 3D bioprinted scaffolding hydrogels to encompass the cellular material in a nutrient and support medium (Calcutt et al., 2021; Jose et al., 2016). In both plants and animals, cells are encompassed by an extracellular matrix, which is an intricate network of cell-material interactions necessary for survival and proliferation of cells through its support and regulation of cellular functions (Durand-Smet et al., 2014; Przekora et al., 2021; Bao et al., 2018). The 3D cellular microenvironment in animals and plants provides significant sources of developmental and biophysical cues Bao et al., 2018; Wang et al., 2016; Miyashima et al., 2019; Schiefelbein et al., 1997; Van Norman et al., 2013). So, when optimizing 3D bioprinting protocols, it is imperative to critically evaluate the selected hydrogel in the bioink so as to avoid creating a harmful cellular microenvironment when bioprinting. There are a wide variety of hydrogels with varying properties such as stiffness, source of derivation, viscosity, temperature or pH range of gelation, nanoporosity, and crosslinking requirements which may all affect bioprinted cells differently since scaffold stiffness and other physical properties can affect the cell viability and shape (Bao et al., 2018; Guimaraes et al., 2022; Lopez-Marcial et al., 2018; Wang et al., 2016). Agarose and Alginate, for example, are naturally-derived hydrogel, or scaffolding, options that are not biodegradable or bioactive (Bao et al., 2018; Guimaraes et al., 2022; Jovic et al., 2019). Alginate hydrogel forms by binding with cations such as calcium while Agarose hydrogel self-gelates as it cools below 37 C (Bao et al., 2018; Jovic et al., 2019). Alginate hydrogels have an elasticity ranging from 10^2 to 10^4 Pa and a lower viscosity than agarose, but alginate hydrogel stiffness can be modified by adjusting its polymer composition, ionic composition, or degree of crosslinking (Wang et al., 2016; Jovic et al., 2019). In mammalian studies, agarose and alginate have both promoted cell proliferation and have been utilized successfully with 3D bioprinting applications (Lopez-Marcial et al., 2018). Pluronic, on the other hand, has been used much more extensively in 3D printing but is not optimally stable in aqueous solutions (Lopez-Marcial et al., 2018; Muller et al., 2015). Compared to pluronic, 2% agarose-based bioink has had a higher viscosity and lower shear rate in comparative studies (Lopez-Marcial et al., 2018). However, pluronic hydrogel can achieve linear lines or patterns in prints whereas agarose or alginate-based hydrogels typically spread beyond the initial print area (Lopez-Marcial et al., 2018). So, hydrogels vary in significant ways and should be chosen and optimized carefully for the desired application. Additional important differences between the hydrogel characteristics include cell-induced material degradation, adhesion, cell differentiation, proliferation, gene expression, protein translation, responsiveness to stimuli, and cellular potency (Wang et al., 2016). Recently, researchers have studied numerous approaches to optimize the bioprinting process (Jose et al., 2016). However, their optimization techniques typically focused on printing parameters, such as flow rate, speed and nozzle size, with a less comprehensive evaluation of various scaffold material properties (Guimaraes et al, 2022). We have previously optimized cell viability and microcallus formation in a 3D bioprinting popeline using *Arabidopsis* root, shoot, and Soybean embryo protoplasts (Van den Broeck et al., 2022). This study aims to achieve further optimization (**Fig. 1**) of the bioprinting process for replicable and repeatable cell viabilities, through the evaluation of more parameters than were previously tested such as a range of extrusion pressure values, needle gauges, and hydrogel typesusing 3D bioprinted BY-2 cells.

**Fig. 1.**
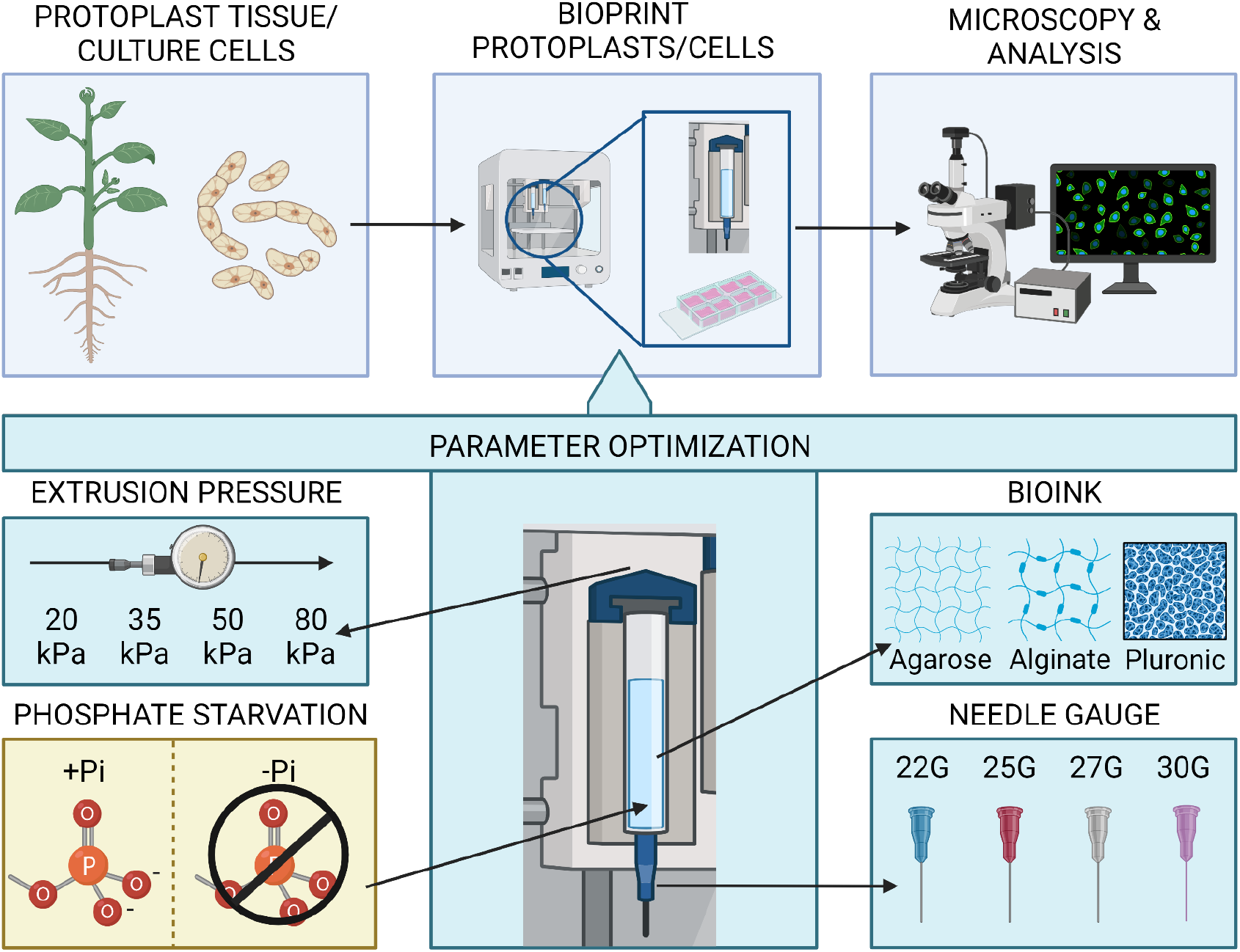
Schematic representation of the 3D bioprinting workflow and parameter optimization. Cells from the Tobacco BY-2 cell culture line or root protoplasts from Arabidopsis seedlings were bioprinted in 8-well chamber slides, according to each tested parameter or condition, then imaged and analyzed to quantify cell viability. The tested parameters included: extrusion pressure (20, 30, 50 or 80 kPa), bioink hydrogel composition (agarose, alginate, and pluronic), and needle gauge (22G, 25G, 27G, or 30G). Finally, 3D bioprinting feasibility of detecting cellular responses to changes in the microenvironment at an optimal set of parameters was quantified using phosphate-starved or phosphate-replete bioink.

## Results

### Scaffold optimization of 3D bioprinted plant cells

We investigated how three hydrogel scaffolds: low melting (LM) agarose (agar), alginate, and pluronic, affected cell viability of BY-2 cells due to their differences in composition. BY-2 cells were bioprinted in each scaffold using a 30G Cellink needle and imaged on the same day after bioprinting (Day 0) (Fig.2a-c). Then, cell viability was calculated of cells bioprinted in either scaffold (Fig. 2d). Cells bioprinted in Agar had a lower cell viability than cells bioprinted in either alginate or pluronic. To identify whether each scaffold had similar effects on cell viability using different bioprinting parameters, BY-2 cells were again bioprinted in each scaffolding material but using a 27G Cellink needle (Fig. 2e-g) instead of a 30G needle and were imaged 2 days later instead of immediately after bioprinting. Interestingly, cell viability was highest using Agar while cell viability was lower using either Alginate or Pluronic with these parameters (Fig. 2h). These differences in cell viability revealed that other factors such as needle diameter may play a larger role in influencing cell viability, particularly in combination with Agar as a scaffold.

**Figure 2.**
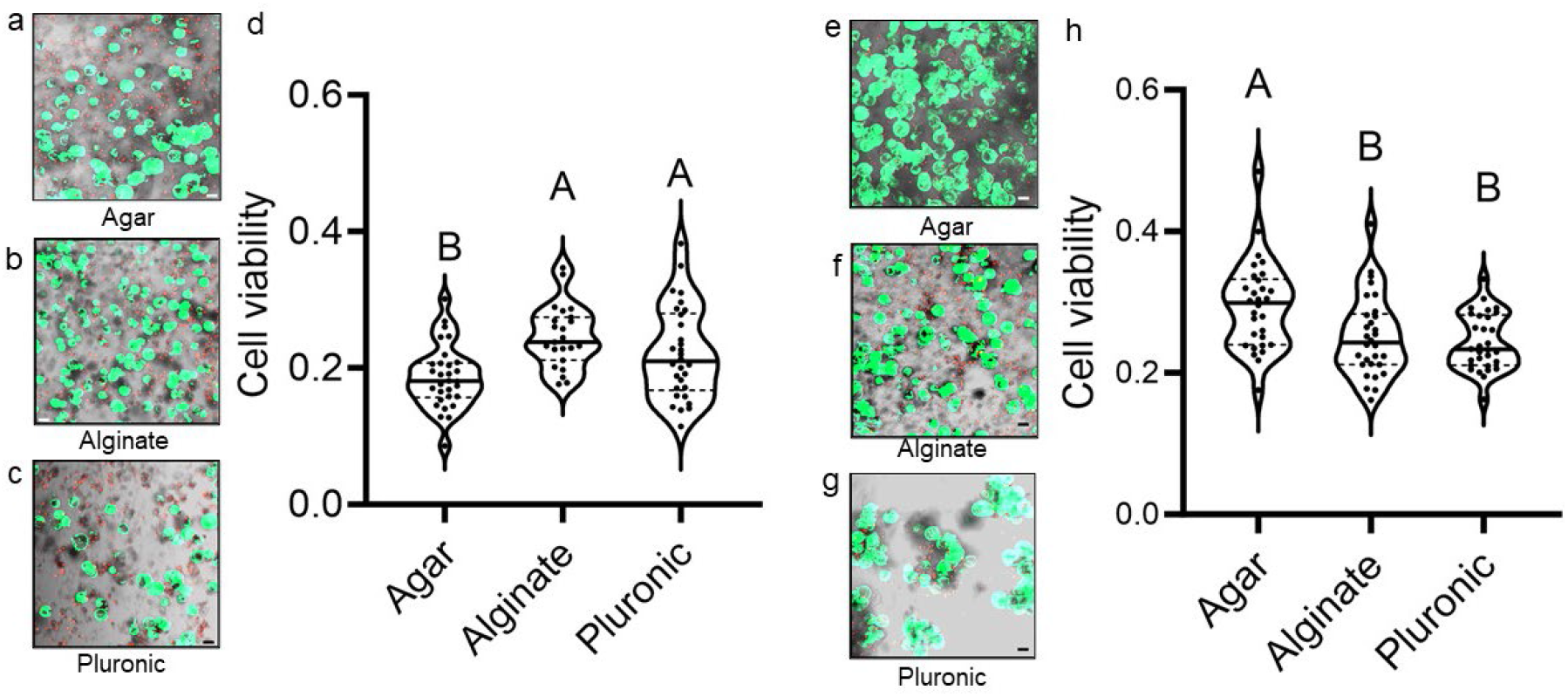
Cell viability of BY-2 cells bioprinted in LM Agarose (Agar), Alginate, and Pluronic scaffolding with either a 30G needle or 27G needle. (**a-c**) Red cells stained with propidium iodide (P.I). indicate dead cells and Green cells stained with Fluorescein Diacetate (FDA) indicate living cells. (**a**) Representative image of BY-2 cells bioprinted in low-melting (LM) agarose. Scale = 50 μm (**b**) Representative image of BY-2 cells bioprinted in alginate. Scale = 50 μm (**c**) Representative image of BY-2 cells bioprinted in pluronic. Scale = 50 μm (**d**) Cell viability of Agar, Alginate, and Pluronic bioprinted with a 30G needle and imaged afterwards in 3 biological replicates. Statistical analysis was performed using a Student’s T Test. (**e-h**) Cell viability of BY-2 cells bioprinted in Agar, Alginate, and Pluronic scaffolding with a 27G needle. (**e-g**) Red cells stained with Propidium Iodide (P.I.) indicate dead cells and Green cells stained with FDA indicate living cells. (**e**) Representative image of BY-2 cells bioprinted in low-melting (LM) agarose. Scale = 50 μm (**g**) Representative image of BY-2 cells bioprinted in alginate. Scale = 50 μm (**g**) Representative image of BY-2 cells bioprinted in pluronic. Scale = 50 μm (**h**) Cell viability of Agar, Alginate, and Pluronic bioprinted with a 27G needle and imaged 2 days afterwards in 3 biological replicates. Statistical analysis was performed using a Student’s T Test.

To identify which additional parameters may be influencing cell viability, BY-2 cells were then bioprinted in an array of needle diameters and extrusion pressures. Since bioprinting with agarose ultimately resulted in the highest cell viability than did Alginate or Pluronic, BY-2 cells were bioprinted using the Agar scaffold in each diameter and pressure combination. Specifically, BY-2 cells were bioprinted at pressure values of either 20 kPa, 25 kPa, 50 kPa, or 80 kPa with needle diameters of either 22G, 25G, 27G, or 30G (Fig. 3a-c). Cell viability was then measured in each pressure-needle combination (Fig. 3d). Using the 22G needle at 20 kPa and 35 kPa, the cell viability was low. Similarly, using the 30G needle at 50 kPa and 80 kPa, cell viability was also low. These data suggest that using needles with a small diameter (30G) at high pressures or large diameter (22G) at low pressures yield poor cell viabilities. The 25G needle yielded a higher cell viability at 20 kPa than at 35 kPa. This suggests that a relatively large diameter yields an improved cell viability at lower pressures. Moreover, the 27G needle yielded a similar cell viability at 35 kPa and at 20 kPa. The 30G needle also yielded a similar cell viability at 35 kPa and at 20 kPa. This suggests that cell viability tolerates a wider range of pressures using relatively small needle diameters. The 27G needle yielded a higher average cell viability at 35 kPa (0.47) than at 20 kPa (0.41) while the average cell viability yielded by the 30G needle had a smaller difference between 20 kPa (0.39) and 35 kPa (0.42). These data suggest that the smaller needle diameters tested (27G and 30G) yield more consistent cell viability results at a pressure range of 20-35 kPa. The 30G needle yielded overall more consistent values between 20 kPa and 35 kPa, suggesting these parameters yielded more consistent cell viabilities.

**Figure 3.**
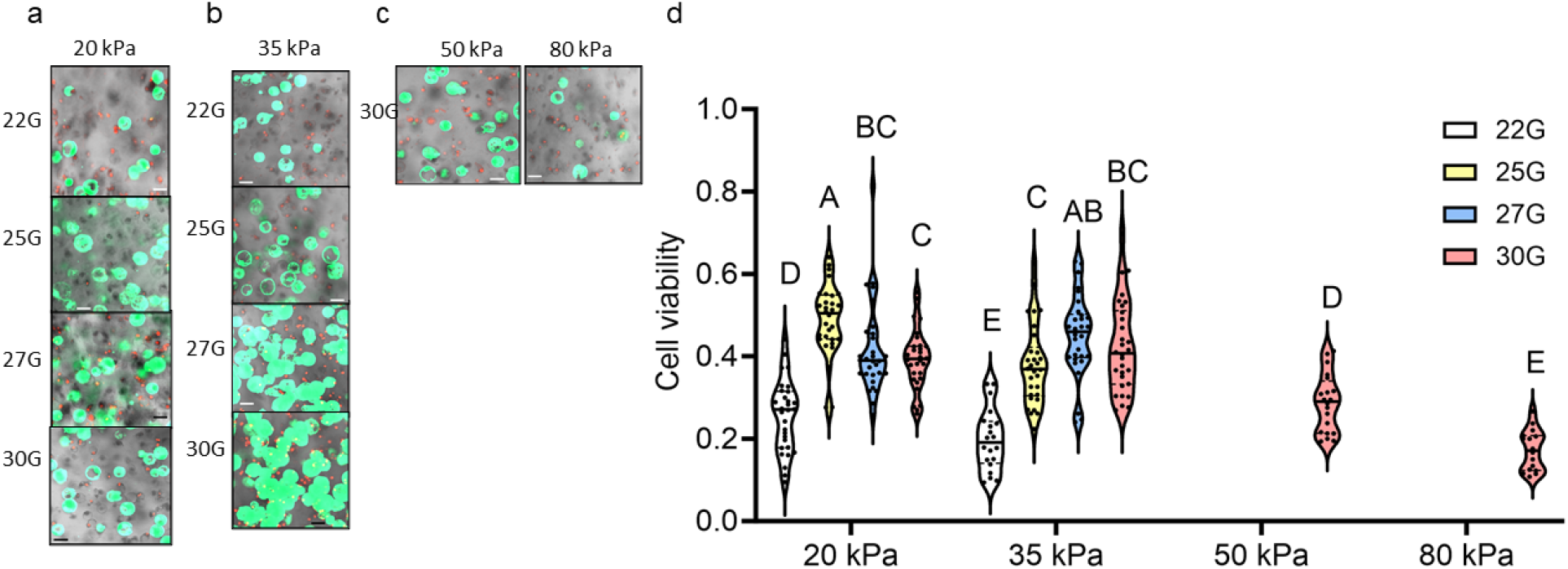
Cell viability of BY-2 cells bioprinted in LM Agarose (agar) using various needles and pressures (**a**) Representative images of BY-2 cells bioprinted using the following parameters: 20kPa at 22G, 25G, 27G, and 30G needle gauges (**b**) Representative images of BY-2 cells bioprinted using the following parameters: 35kPa at 22G, 25G, 27G, and 30G needle gauges. (**c**) Representative images of BY-2 cells bioprinted using the following parameters: 30G needle gauge at either 50kPa and 80 kPa. (**a-c**) Red cells stained with Propidium Iodide (P.I.) indicate dead cells and Green cells stained with Fluorescein Diacetate (FDA) indicate living cells. Imaging was performed immediately after bioprinting. Scale = 50 μm (**d**) Cell viability of BY-2 bioprinted cells using each parameter combination: 22G, 20 kPa (biological replicates n=3); 22G, 35 kPa (n=3); 25G, 20 kPa (n=3); 25G, 25 kPa (n=3); 27G, 20 kPa (n=3); 27G, 35 kPa (n=3); 30G, 20 kPa (n=3); 30G, 35 kPa (n=3); 30G, 50 kPa (n=2); and, 30G, 80 kPa (n=2). Statistical analysis was performed using Student’s T Test.

### 3D bioprinting to simulate and study in field conditions

The identified optimal bioprinting parameters, a 30G needle at 20 kPa using the Agar scaffold, were then applied to a stress condition, phosphate starvation, to investigate whether 3D bioprinting could be an effective system for identifying cell-microenvironment responses. First, root responses such as meristematic cell division and root length were quantified in whole roots of seedlings grown under either Pi-replete (+Pi) at 1.2 mM KH_2_PO_4_ or Pi-starved (-Pi) conditions at 0 mM KH_2_PO_4_.

In roots, the meristematic zone was measured as the distance between the Quiescent Center (QC) and the first elongated cortex and epidermal cells (Verbelen et al., 2006) at 7 days (7d) after sowing on either +Pi (Fig. 4a) or -Pi (Fig. 4b) media and at 10 days (10d) after sowing on either +Pi (Fig. 4c) or -Pi (Fig. 4d) media. The amount of cells present within the meristematic zone of each root was counted as well. Collectively, the length of the meristematic zone and the number of cortical cells in the meristematic zone provide insight into whether primary root growth, as mediated by cell division, is inhibited over time by Pi starvation. At 7d and 10d, Pi starved root meristems were smaller in length (Fig. 4e) and had fewer meristematic cortex cells (Fig. 4f). Between 7 and 10 days, the length of the meristematic zone and number of cortex cells increased in Pi-replete roots, suggesting continued cell division through this time frame. In Pi-starved cells, however, the length of the meristematic zone and number of cortex cells remained similar in Pi-starved cells between 7 and 10 days. This suggests that cell division was inhibited by Pi starvation after 7 days. Finally, root length from 3 to 14 days after sowing on +Pi or -Pi media was measured to determine whether a Pi-starvation induced inhibition of cell division affected primary root growth (Fig. 4g). Pi-starved root growth was consistently inhibited compared to Pi-replete roots within 8-14 days after sowing. This suggests that Pi starvation inhibits root growth likely by inhibiting cell division (Khan et al., 2023).

**Figure 4.**
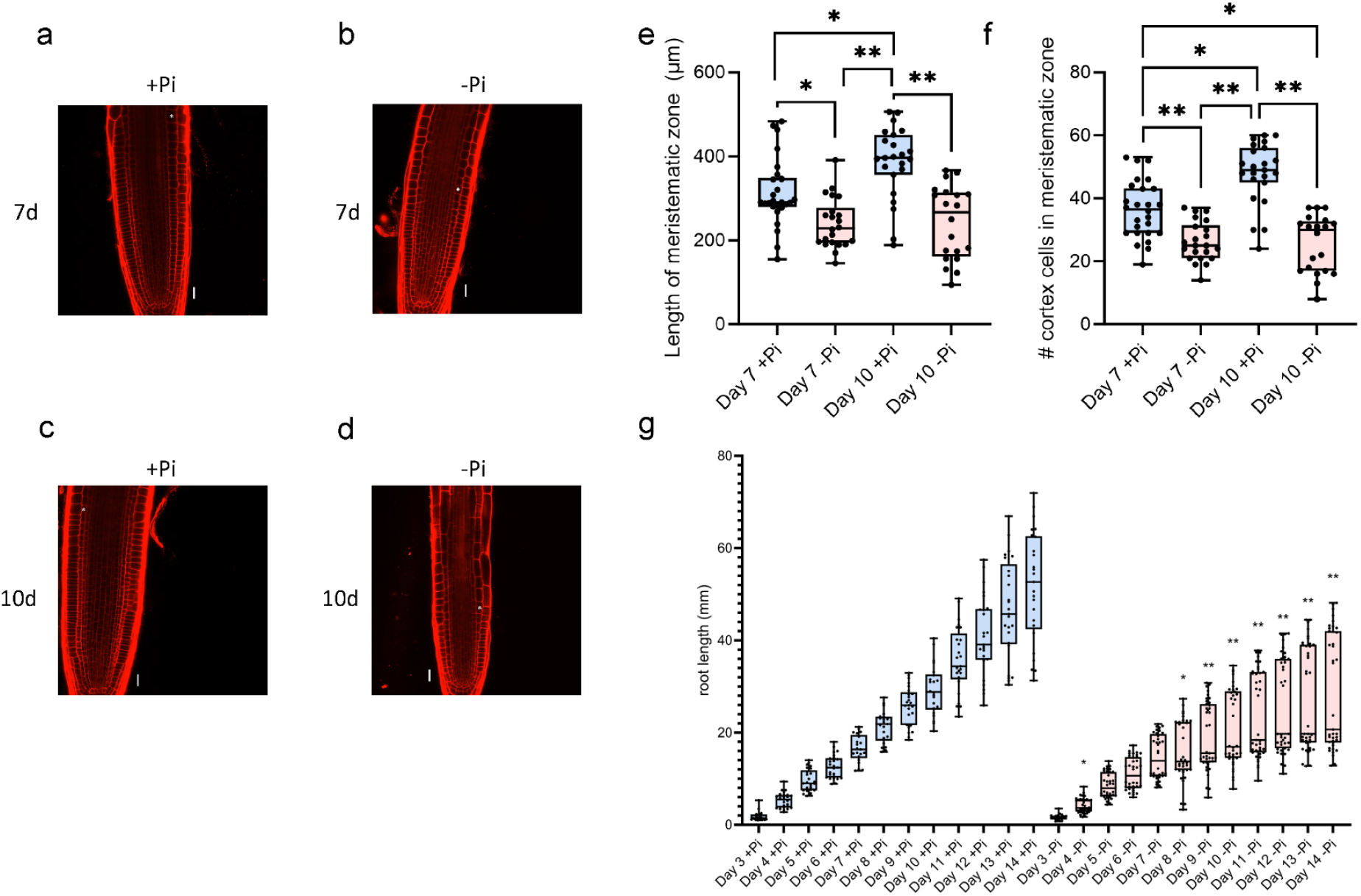
Root meristem division and growth responses to Pi starvation. (**a**) Representative image of +Pi root apical meristem at 7 days after sowing. Scale = 50 μm (**b**) Representative image of -Pi root apical meristem at 7 days after sowing. Asterisks denote end of meristematic zone. Scale = 20 μm (**c**) Representative image of +Pi root apical meristem at 10 days after sowing. Asterisks denote the end of the meristematic zone. Scale = 50 μm (**d**) Representative image of - Pi root apical meristem at 10 days after sowing. Asterisks denote the end of the meristematic zone. Scale = 20 μm (**e**) Size of meristematic zone consisting of stem cell niche and transit amplifying cells. Blue indicates roots grown in +Pi media and red indicates roots grown in -Pi media. The size of the meristematic zone was measured at both 7 and 10 days after sowing. Statistical analysis is a Steel-Dwass test method. * denotes p < 0.05 and ** denotes p < 0.0001. (**f**) Number of cortex cells within the meristematic zone. Blue indicates roots grown in +Pi media and red indicates roots grown in -Pi media. The size of the meristematic zone was measured at both 7 and 10 days after sowing. Statistical analysis is a Steel-Dwass test method. * denotes p < 0.05 and ** denotes p < 0.0001. (**g**) Root length of roots grown in either +Pi media (blue) or -Pi media (red) and root length was measured each day after day 3 of sowing until 14 days after sowing. Statistical analysis is a pairwise Student’s T test comparing root length between +Pi and -Pi at each time point and *p<0.01 and **p<0.0001.

To evaluate whether 3D bioprinting yielded similar results, *Arabidopsis thaliana* protoplasts from 7 day old roots were then bioprinted in either Arabidopsis agarose-based bioink (+Pi) (Van den Broeck et al., 2021) or -Pi bioink. Bioprinted cells were then imaged immediately afterwards and each day up to 3 days and at 7 and 10 days after bioprinting to identify whether root responses and bioprinted cell responses to +Pi and -Pi conditions were similar at both early and late time points (Fig. 5a). There were little differences in cell viability between +Pi and -Pi cells at each day up to 3 days after bioprinting, suggesting that cells were still viable despite the Pi-starved environment (Fig. 5b). So, *Arabidopsis* cells were then bioprinted and cell viability was determined at 7 and 10 days after bioprinting. At both 7 and 10 days, cell viability of Pi-starved bioprinted cells was lower than that of Pi-replete cells, suggesting that bioprinted cells are adversely affected by Pi-starvation at 7 days and afterwards (Fig. 5c). Similarly, root cell division and root inhibition was reduced at 7 days after sowing. These data suggest that both bioprinted cells and cells within whole plants respond similarly to Pi-starvation. Therefore, 3D bioprinting is a useful system for studying cellular microenvironment interactions at the identified 3D bioprinting parameters.

**Figure 5.**
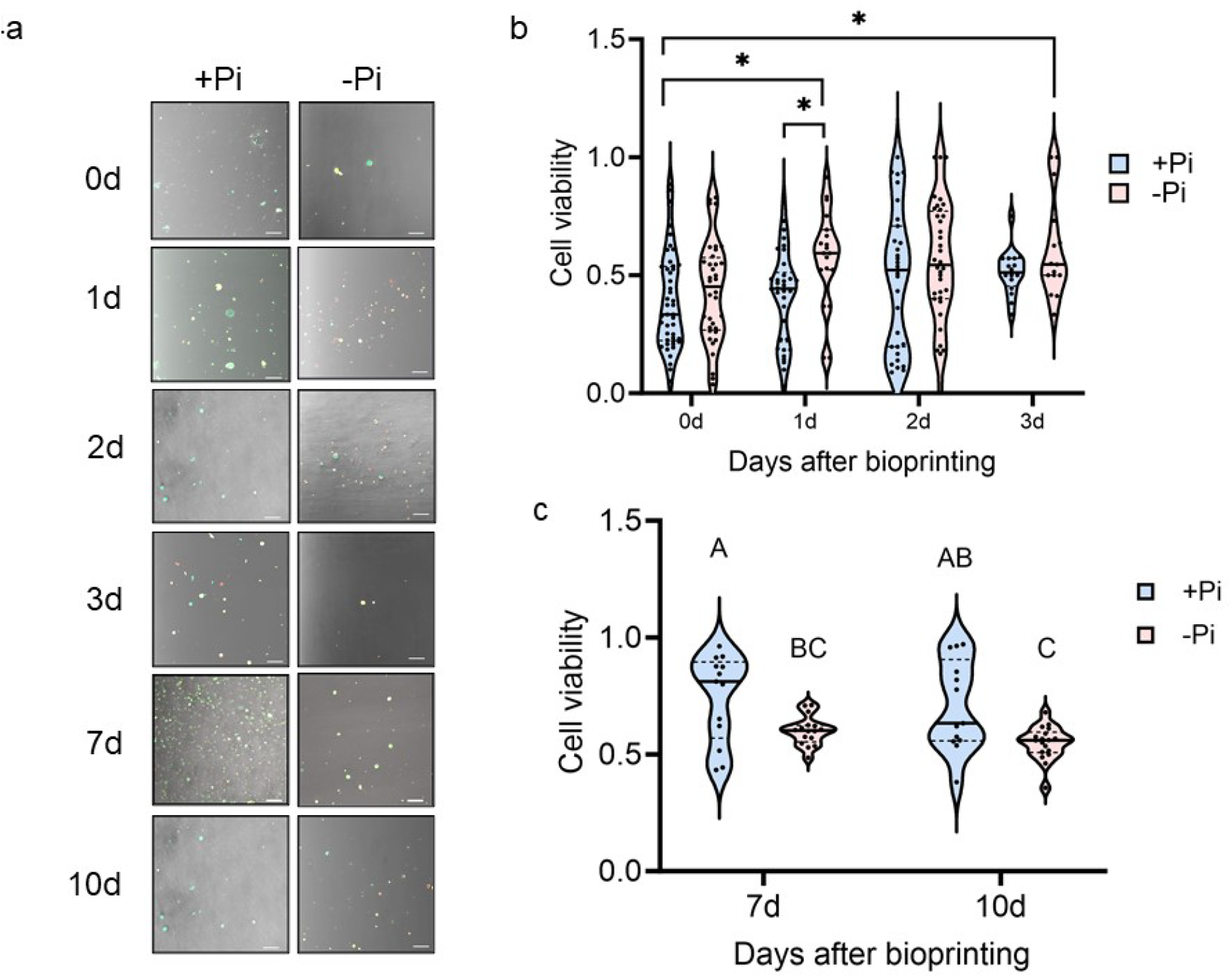
3D Bioprinted cell viability under phosphate-starved and phosphate-replete conditions. (**a**) Representative images of Col-0 *Arabidopsis* root cells under phosphate-replete (+Pi) and phosphate-starved (-Pi) conditions at 0d, 1d, 2d, 3d, 7d, and 10d after bioprinting in either +Pi or -Pi bioink. (**b**) Cell viability results between 0d and 3d after bioprinting Col-0 root cells in either +Pi or -Pi bioink. Statistical analysis was performed using the Steel-Dwas method of pairwise comparisons (**c**) Cell viability results at 7d and 10d after bioprinting Col-0 root cells in either +Pi or -Pi bioink. Statistical analysis was performed using the ANOVA/Tukey HSD method.

## Discussion

3D bioprinting is a promising technology in plant biology for applications such as tissue engineering or cellular microenvironment interaction investigations (Seidel et al., 2017; Mehrotra et al., 2020). Optimizing 3D bioprinting parameters is a necessary step for successfully utilizing this technology in any applications. In past studies, we have optimized protoplasting and bioink formulation protocols in Arabidopsis and soybean (Van den Broeck et al., 2022) . In this study, we have optimized hydrogel and extrusion-based bioprinting parameters including pressure and needle gauge. We have found that agarose hydrogels have a less variable influence on cell viability at different pressure intensities. We have also found that small-diameter needle gauges and pressures between 20-35 kPa are optimal while pressures above 50 kPa dramatically drop cell viabilities in BY-2 cells. Then, the 30G needle at 20 kPa, an optimal set of parameters, was used to bioprint Arabidopsis cells and investigate whether phosphate starvation influenced cell viability over time. As a result of bioprinting with these parameters, we found there was a drop in cell viability after 7 days of phosphate starvation which is consistent with whole root studies. This study presents a significant step towards identifying optimal bioprinting conditions for plant cells especially since most studies include animal or human cells. In future experiments, it will be necessary to determine whether the duration of a bioprinting run influences cell viability and whether a combination of multiple factors (e.g. pressure-gauge-scaffold) combinatorially influence cell viability differently. Composite scaffolds have also been used in mammalian studies, such as agarose-alginate combinations, so further studies should also explore whether, in plant cells, composite scaffolding could even further improve cell viability (Lopez-Marcial et al., 2018). Moreover, there are other considerations outside of cell viability that should be considered such as cell proliferation, cell morphology and membrane integrity, and extra-cellular matrix formation (Guimaraes et al., 2022) to confirm whether bioprinted cells retain physiological characteristics similar to cells *in planta* (Przekora et al., 2019). Since 3D bioprinting has future applications in tissue engineering, then further studies should investigate how cell communication, cell adhesion, and cell detachment occur in 3D bioprinted cells (Przekora et al., 2019; Guimaraes et al., 2022). In particular, this study demonstrates that it is useful to optimize needle gauge and pressure for use with a range of species to give more flexibility in experimental design and applications. Moreover, 3D bioprinting and microfluidics may be technologies useful for studying signal and hormone gradients and their impact on cellular processes (Guimmraes et al., 2022). So, similar assays to this study of evaluating cellular characteristics in response to small parameter changes would be a step towards recapitulating and understanding positional signaling cues on cell identity or behavior. Furthermore, optimizing 3D bioprinting parameters as described in this study will be useful in future studies because it will make clear which parameters would best suit various experimental approaches.

## Materials and Methods

### BY-2 cell culture and isolation

The NT-1 medium for the BY-2 cell culture is made by dissolving 4.3 milliliters (ml) of Murashige & Skoog (MS) Salts, 30 ml of sucrose, 10 ml of B1-Inositol Stock, 3 ml of Miller’s I Stock and 0.02 ml of 2,4 Dichlorophenoxyacetic acid (2,4D) Stock of 10 milligrams per milliliter (mg/ml) into 500 ml of distilled water (H2O) and adjusting the total volume with distilled H2O up to 1000 ml. The 10 mg/ml 2,4-D Stock solution is prepared by melting 0.10 grams (g) of 2,4D into 10 ml of 0.1 Molarity (M) of potassium hydroxide (KOH). B1-Inositol and Miller’s I Stock solutions were made by dissolving 5 g of Myo-Inositol and 0.05 thiamine-hydrochloride into 250 ml of distilled H2O, and 30 g of potassium dihydrogen phosphate (KH2PO4) into 250 ml, respectively; the 2 final volumes are then adjusted to 500 ml with additional distilled H2O.

The BY-2 cell suspension culture is isolated from a 60-milliliter flask of a 7 days old in vitro BY-2 mother culture, which is replenished weekly with fresh nutrient NT-1 medium and kept under constant light in an orbital shaker at a speed of 150 revolutions per minute (rpm) and a temperature of 23o Celsius (C), by transferring 2 ml of cells from the mother culture into a new vial that contains 58 ml of NT-1 medium. The resulting BY-2 cell flask is kept in the same orbital shaker under the same conditions and used for 3D bioprinting on day 3 from the day of its isolation.

### Protoplast isolation

An enzymatic solution must be prepared to break down the cell walls and isolate protoplasts. BY-2 protoplasts are prepared as in (21). Thus, the BY-2 enzymatic solution is made by dissolving 1% (w/v) cellulase (EMD Millipore) and 0.1% (w/v) pectolyase (Sigma-Aldrich) into a 100 ml solution of 99 ml 0.4 M mannitol and 1 mL 500 milli-Molarity (mM) MES. Afterwards, the enzymatic solution is sterilized using a vacuum filter sterilization procedure. For Arabidopsis, the enzyme solution consisted of 1.5% cellulase and 0.1% pectolyase as described in Van den Broeck et al., 2022. Arabidopsis seedlings were grown on MS plates for 7 days at which the roots were cut and placed in a 70-um cell strainer filled with the Arabidopsis enzyme solution. After 1 hour, protoplasts were transferred to a sterile 15 mL conical tube then centrifuged at 500g at 23 C for 5 minutes. The pellet was then resuspended in a 3:1 ratio of Arabidopsis PIM:agarose scaffold before bioprinting (Van den Broeck et al., 2022).

10 ml of the 3 days old BY-2 cell vial is placed into a sterile 50 ml conical tube under the sterile hood and the tube is then centrifuged at a speed of 500 relative centrifugal force (rcf) and a temperature of 23o C for 5 minutes (min). After the tube is spun down, the tube is moved back to the sterile where the supernatant is carefully removed and 7 ml of the enzymatic solution is added to the cell pellet. The tube is meticulously transferred to an orbital shaker and left to stir for an hour at a speed of 85 rpm and a temperature of 23o C. Once the cell wall degradation is completed, the tube is centrifuged at a speed of 500 rcf and a temperature of 23o C for 3 min and the supernatant is carefully removed without disturbance of the protoplast pellet. To do an NT-1 wash that can reduce further effect of the enzymatic solution, 5 ml of NT-1 medium is added to the protoplast pellet and the mixture is slowly pipetted up and down until homogeneity and spun down in the centrifuge at a speed of 500 rcf and a temperature of 23o C for 3 min. The supernatant is thoroughly removed. For LM Agar and Alginate scaffoldings, the protoplast is resuspended in 4 ml of NT-1 medium whereas the resuspension in NT-1 is skipped for Pluronic scaffolding and 4 ml of 20% Pluronic is immediately added to the protoplast pellet for bioprinting.

### Bioink synthesis: low melting (LM) Agar, Alginate, and Pluronic

The experiments in this paper comprise of 3 components which are pressure and nozzle test and the scaffolding test. For the pressure and nozzle test, different combinations of 20, 35, 50 and 80 Kilo Pascal (KPa) pressures and 22, 25, 27 and 30 gauge (G) nozzle sizes are constituted with the utilization of 0.6% of LM Agar to obtain bioink-scaffolding for printing. Similarly, the scaffolding test consisted of 0.6% of LM Agar, 75% of Alginate and 20% of Pluronic. Both LM Agarose and Alginate are kept in the warm before their usage and their corresponding bioinks are prepared with 1 volume of scaffold material and 3 volumes of the protoplast resuspended in NT-1 medium. Pluronic is kept in the sterile hood at room temperature for 7 min as Pluronic is prone to solidify if kept long at temperature that exceeds 4o C and 4 ml of its volume is directly added to the protoplast pellet, as mentioned in the subsection above, to form the bioink. All Arabidopsis Col-0 protoplasts bioprinted in -Pi bioink were resuspended in PIM and agarose scaffold formulated with a homemade B-5 salts recipe in which phosphate was not added. All Arabidopsis protoplasts bioprinted in +Pi bioink were formulated as described in Van den Broeck et al., 2022.

### Bioprinting and construct maintenance

After the addition of the scaffold materials, their bioinks are immediately transferred to their corresponding cartridges and bioprinted to prevent premature solidification of the bioink, which would lead to clotting of the bioink in the bioprinter syringe. Each concentration is bioprinted into an 8-well iBidi slide with 4 constructs in each slide using a Cellink Bioprinter. The bioprinter is calibrated without any temperature adjustment at first and Pluronic bioink is bioprinted soon afterwards. Once the Pluronic bioprinting is completed, the tool of the bioprinter is set to a temperature of 37o C for both LM Agar and Alginate to help maintain the desired viscosity of the bioinks and its printbed at a temperature of 23o C to allow convenient cool down of the bioprinted constructs. For all of the experiments involving LM Agar and Pluronic, 200 microliters (μl) of NT-1 is added to each well of the bioprinted slides, while a crosslinking is done to the Alginate samples, by adding 150 μl of calcium dichloride (CaCl2) to each well and removing it completely after 1 min, before adding 200 μl of NT-1 medium. All the bioprinted samples are kept in the dark in the Percival at a temperature of 23o C.

### Confocal imaging and image analysis

Evaluation of biological characteristics of the bioprinted samples for BY-2 cells is observed using staining, imaging, and digital image analysis. Staining and confocal microscope imaging are done on day 0 for pressure and nozzle test, days 0, 2 & 4 for LM Agar stiffness test and days 0, 1 & 2 for Scaffolding test. Staining is done with fluorescein diacetate (FDA) solution, prepared by mixing 5 mg of FDA into 1 ml of Acetone into a 15 ml conical tube, right before each imaging session to preserve cell viability. 10 μl of 1 M propidium iodide (Pi) solution is diluted into 90 μl of sterile distilled H2O and used for counter staining to determine the nonviable cells. Thus, 10 μl of the diluted Pi solution and 2 μl of the FDA solution are added to each wells of the samples used on imaging days. After staining, the bioprinted constructs of each sample are imaged using a Zeiss LSM 980 confocal microscope and 5-10 3D images with varying z-stacks are generated per each sample. The digital analysis of the confocal images is done as in (Van den Broeck et al., 2022). The results of the image analysis are the viable and nonviable cell counts in csv files used to create a single long data format csv file that is further analyzed and visualized using Python script to quantify the experimental results.

## Acknowledgment

This work was supported by the NSF EAGER grant (MCB #2039285) to TJH and RS. Work by IM was supported by the NSF Postdoctoral Fellowship award (#2305774).

No conflicts of interests are present.

## Notes

### Competing Interest Statement

The authors have declared no competing interest.

